# Developmental shift in testosterone influence on prefrontal emotion control

**DOI:** 10.1101/2022.09.02.505577

**Authors:** Anna Tyborowska, Inge Volman, Hannah C. M. Niermann, Anna L. Dapprich, Sanny Smeekens, Antonius H. N. Cillessen, Ivan Toni, Karin Roelofs

**Author notes:** equal author contribution (Ivan Toni, Karin Roelofs). **Correspondence:** Anna Tyborowska. **Competing Interests** Authors declare that they have no competing interests. **Data and Materials Availability** Data is available upon request via the Donders Repository (https://data.donders.ru.nl/). Data can be provided by the Donders Institute for Brain, Cognition, and Behaviour pending scientific review and a completed data transfer agreement. Requests for the data should be submitted to AT or KR. **Ethical Approval** Written informed consent was obtained from both parents and participants. The study was approved by the local ethics committee (CMO region Arnhem-Nijmegen, The Netherlands).

## Abstract

A paradox of testosterone effects is seen in adolescents vs. adults in social emotional approach-avoidance behavior. During adolescence, high testosterone levels are associated with increased anterior prefrontal (aPFC) involvement in emotion control, whereas during adulthood this neuro-endocrine relation is reversed. Rodent work shows that, during puberty, testosterone transitions from a neuro-developmental to a social-sexual activating hormone. In this study, we explored whether this functional transition is also present in human adolescents and young adults. Using a prospective longitudinal design, we investigated the role of testosterone on neural control of social emotional behavior during the transitions from middle to late adolescence and into young adulthood. Seventy-one individuals (tested at ages 14, 17, and 20 years) performed an fMRI-adapted approach-avoidance (AA) task involving automatic and controlled actions in response to social emotional stimuli. In line with predictions from animal models, the effect of testosterone on aPFC engagement decreased between middle and late adolescence, and shifted into an activational role by young adulthood - impeding neural control of emotions. This change in testosterone function was accompanied by increased testosterone-modulated amygdala reactivity. These findings qualify the testosterone-dependent maturation of the prefrontal-amygdala circuit supporting emotion control during the transition from middle adolescence into young adulthood.

## Introduction

The gonadal hormone testosterone plays a pivotal role in social emotional behavior by modulating the control of approach and avoidance actions (1–3). However, the function of testosterone changes during puberty (4, 5). For instance, higher levels of pubertal testosterone have been associated with *increased* anterior prefrontal (aPFC) emotion control in mid adolescence (1), whereas in adult populations, higher levels of testosterone have been associated with *reduced* aPFC emotion control (2, 6). Here, we use a prospective longitudinal design to understand how the human prefrontal cortex matures into adult-like regulation of social-emotional behavior and most critically, to test the changing role of testosterone in this maturation.

The paradox of seemingly opposing testosterone effects between adolescents and adults is particularly relevant for understanding the development of social approach-avoidance behavior. Studies in humans confirm that pubertal testosterone modulates executive control and reward processing networks (7–12). Importantly, testosterone has been linked to the maturation of cognitive control over social-emotional actions, a fundamental adult skill refined during puberty. During mid-adolescence, relatively more mature adolescents, as indexed by higher levels of testosterone, implement emotion control by relying on the aPFC. Interestingly, at the same age, less mature individuals rely on pulvino-amygdala interactions to control their emotional action tendencies -a precursor of the aPFC-amygdala circuit typically used by adults (1). In contrast, adults’ testosterone in adults is associated with dominance, status-seeking and aggressive behavior (13, 14). In line with this notion, adults with high testosterone levels rely less on the aPFC and have reduced aPFC-amygdala coupling during emotion control (2).

Animal work specifies that the function of testosterone during adolescence shifts from a developmental agent of neural organization to a hormone transiently activating socio-sexual behavioral and neural responses (4). Pubertal testosterone is critical for inducing long-lasting system changes and programing adult social behavior (15). For instance, animals that are castrated after infancy will display typical socio-sexual and territorial (dominance) behavior as adults only if they are given testosterone-replacement treatment in the pre- and mid-adolescent phase, and not later in the post-adolescent phase (16, 17). Similar effects have been shown for adolescent maturation of social learning and adaptive sexual approach behavior (17, 18). This body of work indicates that the organizational–activation function of gonadal hormones applies particularly to programing and regulation of context dependent social approach-avoidance motivation during adolescence (15). However, it remains to be seen whether testosterone shifts from a neuro-developmental to a social-sexual activating hormone during human adolescence. Here, we tested this hypothesis in a longitudinal study on emotion control using a task that assesses approach-avoidance responses in humans. Knowing whether and how testosterone changes function during adolescence is necessary for understanding adaptive development, as well as maladaptive maturational processes, which may lead to psychopathology, such as anxiety or aggression-related disorders.

Using new data acquired in the same individuals tested in (1), we tested whether testosterone levels in middle and late adolescence and young adulthood predicted the engagement of the aPFC during emotion action control. Middle and late adolescence marks two critical developmental periods. At age 14 (mid adolescence), testosterone is still playing a largely developmental role, with testosterone levels differing widely between individuals (19, 20). At age 17 (late adolescence), testosterone has less influence as a pubertal hormone, and has reached peak levels for girls (21). Hormone levels will continue to increase for boys until their early twenties, highlighting the developmental offset, as well as the sex differences that emerge at this age (8, 21).

We used the approach-avoidance (AA) task, an experimental task that captures behavioral and neural components of emotion action control in adolescents and adults (1, 2, 22, 23). During this task, participants need to evaluate the emotional expression (happy, angry) of faces and respond by pulling a joystick toward themselves (approach) or pushing it away (avoidance). Automatic ‘affect-congruent’ stimulus response mappings (i.e. approach-happy and avoid-angry faces) are contrasted with ‘affect-incongruent’ conditions requiring the participant to override these emotion tendencies and select an alternative goal-directed action (approach-angry and avoid-happy). The aPFC is important for this type of emotion control (defined as the above-described comparison) and plays a role in down-regulating amygdala reactivity (1, 6, 24, 25).

Following the organizational–activation hypothesis of gonadal hormones in rodents, we hypothesized that the modulating effect of testosterone on aPFC emotion control will relatively decrease between middle and late adolescence and switch to negative modulation by young adulthood (Figure 1A). Specifically, we examined this by investigating the change of the congruency effect over time, that is, the difference between affect-incongruent (happy-avoid, angry-approach) and affect-congruent (happy-approach, angry-avoid) trials, in association with testosterone at each age. We focused on the aPFC, amygdala, and pulvinar, given their previously demonstrated involvement in social emotional control (1).

**Fig. 1.**
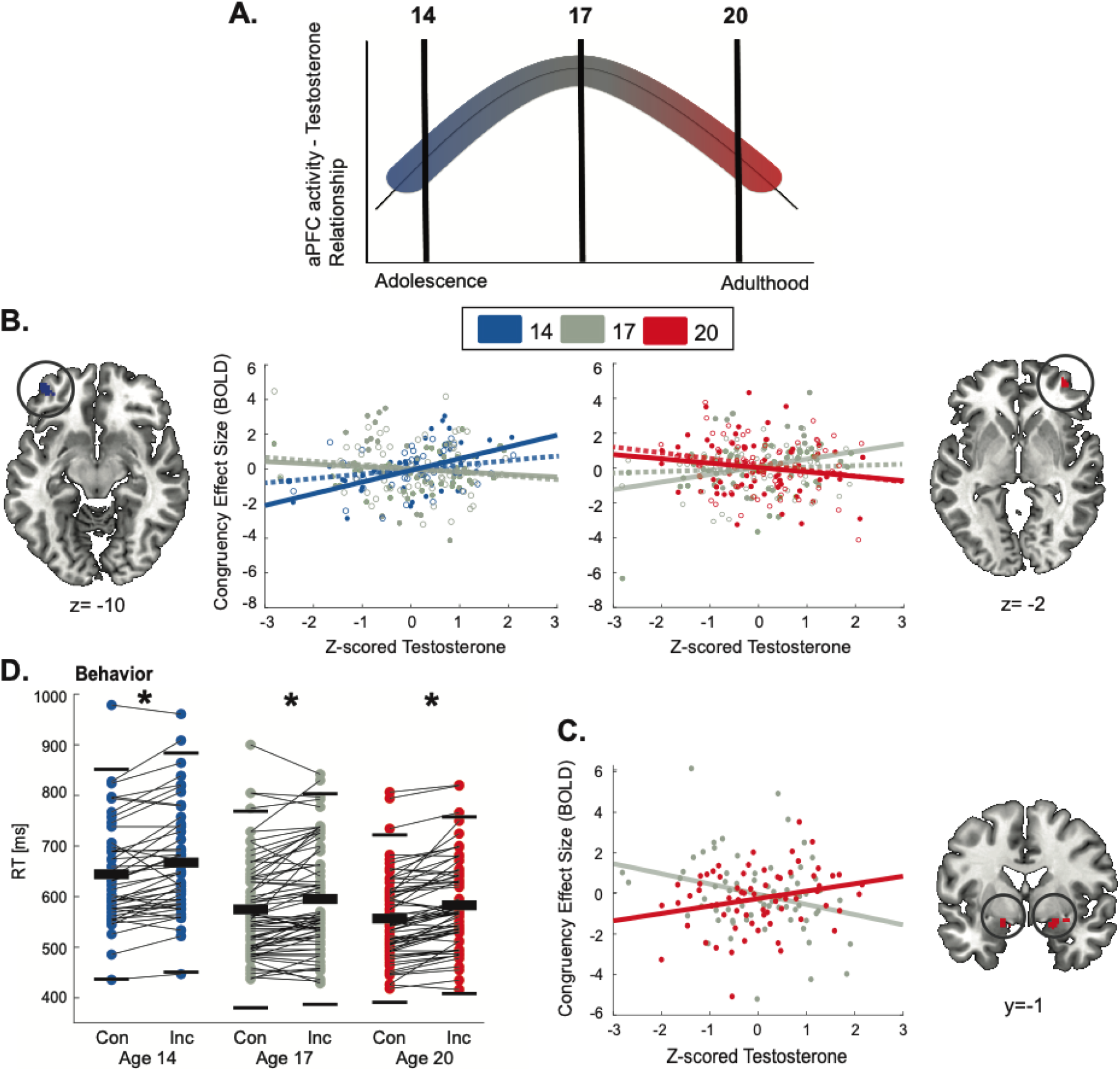
Testosterone modulation of emotional control (congruency effect - incongruent vs. congruent trials) across time. **A)**. Hypothesized transition of testosterone-modulation effects on aPFC function from adolescence to young adulthood **B)**. Testosterone modulation of aPFC activity during emotional control – aPFC decreased from age 14 to age 17 and shifted to a negative association by age 20 **C)**. Amygdala activity increase at age 20 compared to age 17. Scatterplots show the mean-adjusted BOLD congruency effects (incongruent > congruent) for age 14 (blue), age 17 (grey) and age 20 (red). Subthreshold effects from the opposing hemisphere (cluster-uncorrected p=.001) are represented by dotted lines and open circles. For visualization purposes activation clusters are presented at a cluster-uncorrected threshold of p < .001. **D)**. Congruency effect [Incongruent (Inc) > Congruent (Con) trials] for reaction times on the AA task at age 14, 17, and 20. Markers represent the average congruency effect per participant. Thick black bars represent the group mean and thin lines +/- 2SD.

## Materials and Methods

### Participants

Children from the Nijmegen Longitudinal Study on Child and Infant Development (NLS; *n* = 116) were approached to take part in this imaging study. Participants were recruited from a range of social economic backgrounds, which is representative of the Dutch population of families with children in the same age range. Of note, the restricted variation of social economic status (SES) in the Netherlands is likely less than in other countries. For more information on the NLS sample and SES see (26, 27). Functional magnetic resonance imaging (fMRI) was obtained at ages 14, 17, and 20 years (Table 1). At the first time-point (TP14), data was obtained from 49 participants, at the second time-point from 73 participants (TP17), and at the third time-point from 69 participants (TP20) who had no counter-indications for undergoing MRI (metal objects in upper body, implants, claustrophobia, etc.). Two participants at TP14 and two at TP17 were excluded due to poor fMRI data quality. (f)MRI data was not collected for one participant at TP20 who stopped after the training phase of the task. This resulted in a total sample of 47 participants (21 males) at TP14, 71 participants (40 males) at TP17, and 68 participants (36 males) at TP20. All participants had normal or corrected-to-normal vision and no history of neurological illness (as indicated by parent/guardian report). Written informed consent was obtained from both parents and participants. The study was approved by the local ethics committee (CMO region Arnhem-Nijmegen, The Netherlands).

**Table 1.** Demographic information: Testosterone levels, PDS scores, age per sex and time-point.

|  | Testosterone s1<br>(pg/ml) | Testosterone s2<br>(pg/ml) | PDS score | Age |
| --- | --- | --- | --- | --- |
| *Age 14 (n=47)* |  |  |  |  |
| Males | 39.4 (31.5) | 42.2 (24.8) | 2.4 (0.6) | 14.55 (0.10) |
| Females | 10.7 (6.3) | 11.7 (9.3) | 2.6 (1.0) | 14.61 (0.18) |
| *Age 17 (n=71)* |  |  |  |  |
| Males | 116.9 (62.8) | 129.0 (74.2) | 3.4 (0.5) | 17.22 (0.16) |
| Females | 28.0 (16.0) | 25.6 (14.4) | 3.5 (0.4) | 17.09 (0.16) |
| *Age 20 (n=68)* |  |  |  |  |
| Males | 139.6 (69.1) | 157.9 (68.9) | N.A. | 20.61 (0.13) |
| Females | 40.1 (32.5) | 51.6 (34.2) | N.A. | 20.62 (0.21) |

Values represent the mean (SD) across participants. Testosterone s1 and s2 represent the first and second (i.e. before and after the task) salivary measurement respectively. Testosterone s1 levels were used for all subsequent analyses. PDS, Pubertal Development Scale; N.A., not applicable.

### Experimental task

In the fMRI adapted AA task, participants responded with a joystick to visually presented emotional faces (happy, angry) by pushing them toward themselves (approach) or away from themselves (avoid). Two block types/response mappings (affect congruent, affect incongruent) lead to four affect x response combinations: happy– approach, angry–avoid (affect congruent) and happy–avoid, angry–approach (affect incongruent).

The task consisted of 16 blocks, 12 trials each. Each block was followed by a baseline interval (21–24 s). The participant was instructed on the required response mapping at the start of each block. The mapping changed after each block, with the order counterbalanced across participants. The affective expressions and the sex of the faces were evenly and pseudo-randomly distributed in each block – there were no more than three consecutive presentations of each. At the start of the task, participants completed a training phase, which consisted of four blocks, eight trails each and the same affect x response combinations. The facial stimuli used in the training differed from the ones used in the experimental phase of the task.

Valid responses were defined as at least 65% of joystick displacement along the sagittal plane and delivered within 3 s following stimulus presentation. Invalid responses were signaled for 1 s with visual feedback indicating “you did not move your joystick far enough.” Once the response was completed, the joystick was returned to its starting position (defined as the central area covering 15% along the sagittal plane) before the end of the inter-trial interval (ITI; 2–4 s). If participants did not do this, they received visual feedback stating “return the joystick to the starting position”. The ITI was repeated after the joystick was returned to its correct position.

### Materials and apparatus

The fMRI data were acquired on a Siemens 3 Tesla MAGNETOM Trio and PRISMA MRI scanner (at same testing site) equipped with a 32-channel coil. The same multi-echo echoplanar imaging (EPI) sequence was used for acquisition of functional scans during all time-points (TR = 2190 ms; TP14 TE = 9.3, 20.9, 32, and 44 ms / TP17 TE = 9, 20.6, 32.16, and 43.74 ms / TP20 TE = 9. 20.4, 31.8, and 43.2 ms; flip angle = 90°; 34 transversal slices; 3.3 × 3.3 × 3.0 mm voxels; FOV = 212 mm). The multi echo EPI sequence is a parallel imaging technique for functional images. It enables a significant reduction in the echo train length, which reduces motion artifacts, image distortion and improves BOLD sensitivity - especially in brain regions typically affected by the use of a single short TE. As a result, it also allows for a better coregistration of functional and anatomical data (28). Our data was acquired on two different scanners due to an update of the resonance magnetic system during data collection at the testing site. As a result, at TP17, 45 out of 71 participants were scanned on the PRISMA system. T1 anatomical scans were collected using an MPRAGE sequence (TR = 2300 ms; TE = 3.03 ms; 192 sagittal slices; 1.0 × 1.0 × 1.0 mm voxels; FOV = 256 mm).

The stimuli consisted of faces, taken from several databases (29–32). There were 36 facial models (18 male) with two affective expressions (happy, angry) each, presented in grayscale against a black background, matched for brightness and contrast values. The faces were trimmed to exclude influence from hair and non-facial contours (based on (23)) and were presented at the center of the screen (visual angle 4° x 6°), visible through a mirror above the participant’s head. Push and pull responses were made with an MR-compatible joystick (Fiber Optic Joystick, Current Designs), placed on the abdomen, with a sampling rate of ∼ 550 Hz. Stimuli presentation and acquisition of joystick positions were controlled by a PC running Presentation software version 10.2 (http://www.neurobs.com).

### Procedure

A similar procedure was followed for all time-points (for more information on TP14 see: (1)). During all visits, participants had the option of familiarizing themselves in a dummy scanner (a simulation scanner that lacks the magnetic field of the MRI scanner). During TP14, participants also filled out several questionnaires prior to entering the scanner and completed an additional non-emotional fMRI task before the experiment began. During TP17 and TP20, no other testing occurred before the experimental task. Saliva samples were always collected before participants were positioned in the MRI scanner. Participants were familiarized with the setup of the AA task in a short training session (10 min), immediately followed by the fMRI session (20 min) and an anatomical scan (5 min). After exiting the scanner, a second set of saliva measurements was collected.

### Pubertal development measures

Saliva samples for testosterone and cortisol measures were collected into Salicap (IBL) containers by passive drool of ∼ 2ml and stored at -20 to -24°C, following the procedure used in (1). Testosterone concentration was measured using a competitive chemiluminescence immunoassay (CLIA) with a sensitivity of 0.0025 ng/mL (IBL) and intra-assay and inter-assay coefficients between 10% and 12%. Cortisol concentration was measured using a commercially available CLIA with high sensitivity of 0.16 ng/mL (IBL) and intra-assay and inter-assay coefficients of <8%.

Participants were instructed to refrain from consuming any food, cigarettes, and drinks (except water) at least ∼ 1 h before the experiment and to refrain from heavy physical exercise. To minimize any possible effects of testosterone level fluctuations, which vary the most in the early morning hours (33, 34), the hormone was sampled in duplicate, 2 hours apart, after 10:00 A.M., resulting in consistent measurements across these two sampling times. Additionally, females were tested outside of the menstruation phase of their cycle. Finally, a self-report measure of pubertal development (secondary sexual characteristics) was collected with the use of the Pubertal Development Scale (PDS; (35)) at TP14 and TP17.

### Behavioral analysis

The behavioral data was preprocessed with Matlab 2012 (MathWorks). Movement onset was reconstructed from the joystick displacement measures on each trial (1, 2). Reaction time (RT) was calculated as the time from picture presentation to movement onset. Trials with no response or a joystick movement in the incorrect direction were not included in further analysis. Trials with an extreme RT (< 150 and > 1500 ms), and a RT > 3SD from the mean were excluded. If a block had an error rate (ER) above chance level, it was excluded in its entirety. RTs were log-transformed to correct for a skewed distribution. Testosterone and cortisol levels taken from first salivary sample (per time-point) were log transformed and standardized per sex.

RT data analysis was carried out in R version 3.6.1 and RStudio version 1.2.5019. A Bayesian mixed-effects model analysis was used to investigate the change in the congruency effect - the difference between affect-incongruent (happy-avoid, angry-approach) and affect-congruent (happy-approach, angry-avoid) trials - over time and in association with testosterone. Data was analyzed at the trial level with Stan (a computational framework for Bayesian analyses (36)), using the brms package (37). The fixed effects of age (TP14, TP17, TP20), valence (happy, angry), response (approach, avoid), testosterone (TP14, TP17, TP20), cortisol (TP14, TP17, TP20) and sex (male, female) were entered into the model, including all underlying lower level interactions and main effects. A random slope was specified for the fixed effects of age, valence, response and testosterone, and their interactions and a random intercept per participant and facial model of the task. An apriori successive difference contrast was defined for the factor age. This contrast tests the differences between the means of the second and first levels (TP17 vs. TP14) and the third and second levels (TP20 vs. TP17)). Estimates were deemed significant if the 95% credible interval (CI; Bayesian confidence interval) did not include zero.

Model convergence was checked with Rhat values (<= 1.01) and caterpillar plots. Weakly informative (neutral), default priors of the brms package (37, 38) that do not substantially influence results were used (no prior information to derive informative priors from was available). Due to convergence issues, not all initially planned random slopes for each fixed effect could be added to the model (i.e. cortisol and gender). As a result, best practice guidelines were followed to ensure a possible maximal random effects structure (39). All possible random covariance terms between random effects were added.

Finally, a quality control analysis was carried out on testosterone levels to check for large differences between s1 and s2 levels, i.e. between samples taken before (s1) and after (s2) the task. A paired-sample *t*-test was conducted on raw testosterone s1 and s2 levels per time-point, separately for girls and boys.

### Functional MRI preprocessing and analysis

Functional data were preprocessed and analyzed using the Matlab toolbox SPM12 [Statistical Parametric Mapping (www.fil.ion.ucl.uk/spm)]. The multi-echo sequence acquired four echoes per volume at every point in time. To control for T1 equilibration effects, the first four volumes of each echo were discarded. The separate echoes were then combined into a (echo-time weighted) single volume. Thorough de-noising of the functional MRI images was performed to remove motion artefacts. First, we used principle components analyses (PCA) to filter out slice-specific noise components based on the shortest echo. Head motion parameters were then estimated using a least-squares approach with six rigid-body transformation parameters (translations, rotations). Next, the time courses of each voxel were realigned to the middle slice (slice 17) to correct for time differences in acquisition. The T1-weighted image was then coregistered to the mean of the functional images and was segmented into grey matter, white matter, and CSF using the “Segment” tool in SPM12. The fMRI time series was normalized and smoothed using a group-specific template based on T1-weighted images. For longitudinal analyses, the relevant fMRI time series were normalized and smoothed using a group-specific template created from T1 images taken at both time-points. Inter-subject registration of the grey matter images was done using diffeomorphic anatomical registration through exponentiated lie algebra (DARTEL; (40)). The fMRI time series was then transformed and resampled at an isotropic voxel size of 2 mm, resulting in spatially normalized, Jacobian scaled, and smoothed (8mm FWHM Gaussian kernel) images. As a following de-noising step, we used ICA-AROMA (independent component analysis-based automatic removal of motion artefacts), run in the FSL environment, to remove noise components in an automated, observer-independent manner (41, 42).

The fMRI time series was analyzed within the general linear model as an event-related design; separately for TP14, TP17, TP20. The vectors containing the onset and duration of each condition (approach-happy, approach-angry, avoid-happy, avoid-angry) were convolved with a canonical hemodynamic response function. Trials that were excluded from the behavioral analysis (misses) as well as instructions or feedback were also modeled. This resulted in six task-related regressors. Additionally, residual head movement-related effects were modeled using original, squared, cubic, first-order, and second-order derivatives of the movement parameters estimated with the spatial realignment procedure (43). The time series of signal intensities from white matter, CSF, and out-of-skull areas were included as additional regressors. The inclusion of these additional regressors helps with image intensity shifts due to hand movement within or near the scanner’s magnetic field (44). The fMRI time series was high-pass filtered with a cutoff of 128 s and corrected for temporal autocorrelation using a first-order autoregressive model.

### Linear mixed effects analyses

Four contrast images of the effects of interest (incongruent: approach-angry, avoid-happy; congruent: approach-happy avoid-angry) were created per subject and entered into Neuropointillist (http://ibic.github.io/neuropointillist) - a toolbox in R that computes voxel-by-voxel statistics. We ran a linear mixed-effects analysis from the nlme package in R version 3.5 with fixed factors Age, Congruency (incongruent, congruent), testosterone, cortisol, scanner, and sex, including the interaction of age x congruency x testosterone x cortisol as well as all lower level interaction terms. A random slope was specified for age and a random intercept per participant. Testosterone and cortisol levels were log-transformed and standardized per sex. Scanner was included to control for the acquisition of data on different machines. Testosterone was considered a covariate of interest. The inclusion of cortisol as a covariate of no-interest is in line with previous neuroimaging studies on the AA task (1, 2) as well as its role in modulating social motivational behavior (23, 45) and was, therefore, included in the model to interact with the primary effects of interest (i.e. the change over time in the testosterone modulation of the congruency effect). An apriori sequential difference contrast was used to compare TP17 vs. TP14 and TP20 vs. TP17. This approach allows testing differential linear developmental trajectories between time-points, as opposed to implementing a higher order (quadratic) function, which poses a risk of over-fitting data to more complex trajectories when there is a maximum of three data points for each individual.

The analysis was run on the whole brain as well as three bilateral apriori regions of interest (ROI) - lateral frontal pole (lateral aPFC) mask from (46), amygdala mask from the Automated Anatomical Labeling (AAL) Atlas (47) and a pulvinar mask from the Automated Tailarach Atlas. The masks were consistent with those used in the previously reported study (1).

The resulting brain images were thresholded at a voxel level (*p*<.001) and cluster (two-sided α = .05, NN = 3) extent using AFNI’s 3dClustSim after estimating the spatial structure of the noise in the dataset using the auto-correlation function (ACF) parameters obtained from the residual timeseries. The cluster extent was computed separately for each ROI (aPFC = 16.8, amygdala = 2.1, pulvinar = 3 voxels) and the whole brain (357 voxels). Anatomical inference was drawn by superimposing the images on a standard SPM single-subject T1 template and subject specific T1 scans in MNI space. Brodmann areas (BA) were assigned based on the MRIcron template (http://people.cas.sc.edu/rorden/mricron/index.html).

### Functional connectivity analyses

We tested whether changes between TP14, TP17 and TP20 in amygdala-prefrontal interregional connectivity are present as a function of pubertal development. We performed a psychophysical interaction (PPI) analysis (48) with the left and right aPFC and right amygdala as seed regions during affect-incongruent versus affect-congruent conditions of the AA task (see fMRI results). Testosterone TP14, TP17, and TP20 levels (log-transformed) were standardized per sex and used as covariates.

For each participant, seed voxels were selected based on a sphere (8mm radius) created around the peak voxel of the group level testosterone-modulated activation for the congruency effect change between TP14 vs. TP17 (MNI coordinates aPFC: -40, 50, - 10) and TP17 vs. TP20 (MNI coordinates aPFC: 38, 50, -2; amygdala: 20, 0, -14). Next, contrast images for the PPI were generated between the seed region time courses and affect-incongruent versus affect-congruent conditions. These participant-specific contrast images (for each time-point) were entered into a linear mixed effects analysis with factors age, testosterone, cortisol, sex, and scanner, including the interaction term age x testosterone x cortisol as well as all lower level interactions, a random slope for age and a random intercept for each participant. Similar to previous analyses, a sequential difference contrast was used to compare TP17 vs. TP14 and TP20 vs. TP17. We performed an ROI analysis on the aPFC timecourses with a bilateral amygdala mask taken from the AAL Atlas (47) and a lateral frontal pole (lateral aPFC) mask from (46) for the amygdala time course.

## Results

### fMRI Results

#### Changes in testosterone modulation of congruency effect

To investigate the developmental trajectory of testosterone on emotion control, we first tested for age differences between TP17 vs. TP14 and TP20 vs. TP17 in the testosterone-modulated congruency effect. We found significant interactions between age x congruency x testosterone in the aPFC for both the TP17 vs. TP14 and TP20 vs. TP17 comparison. While higher testosterone levels were associated with stronger left aPFC activity at TP14 (replicating (1)), this effect decreased at TP17 (MNI coordinates: -40, 50, -10; Figure 1B). The comparison between TP20 vs. TP17 revealed a shift towards the opposite effect - higher testosterone levels at TP20 were related to less right aPFC activity during control of emotional actions (MNI coordinates: 38, 50, -2; Figure 1B). The TP20 vs. TP17 comparison also revealed a significant interaction between age x congruency x testosterone in the bilateral amygdala (MNI coordinates: 20, 0, -14; -16, -2, -12) indicating higher testosterone levels related to more amygdala reactivity at TP20, with the opposite pattern present at TP17 (Figure 1C). Finally, there was a significant testosterone-modulated congruency effect for TP17 vs. TP14 in the bilateral pulvinar (MNI coordinates -16, -28, 12; 24, -24, 12), replicating the negative association previously reported at TP14 (1) and lack thereof at TP17. For all ROI effects see Table 2; for whole-brain effects see Supplementary Information Table S1.

**Table 2.** Region of interest significant clusters and peak activation changes with age during affect incongruent compared to affect congruent trials in the AA task.

| Anatomical region | Side | BA | k | MNI coordinates |  |  | *p* | t |
| --- | --- | --- | --- | --- | --- | --- | --- | --- |
|  |  |  |  | x | y | z |  |  |
| *Congruency effect* |  |  |  |  |  |  |  |  |
| *TP17 vs. TP14* |  |  |  |  |  |  |  |  |
| aPFC | R | 11 | 37 | 24 | 66 | 2 | .0001 | 4.1 |
| *TP20 vs. TP17* |  |  |  |  |  |  |  |  |
| aPFC | L | 11 | 18 | -28 | 58 | -12 | <.0001 | 4.21 |
| Pulvinar | R | - | 87 | 18 | -30 | 4 | <.0001 | 5.78 |
| *Testosterone modulation of congruency effect* |  |  |  |  |  |  |  |  |
| *TP17 vs. TP14* |  |  |  |  |  |  |  |  |
| aPFC | L | 10, 47 | 126 | -40 | 50 | -10 | <.0001 | 4.76 |
| Pulvinar | L | - | 11 | -16 | -28 | 12 | .0001 | 3.85 |
| *TP20 vs. TP17* |  |  |  |  |  |  |  |  |
| aPFC | R | 47 | 23 | 38 | 50 | -2 | .0002 | 3.71 |
| Amygdala | R | 34 | 24 | 20 | 0 | -14 | <.0001 | 4.63 |
|  | L | 34 | 17 | -16 | -2 | -12 | <.0001 | 4.50 |
|  | R | 34 | 7 | 28 | -4 | -14 | .0004 | 3.53 |
| *Testosterone and cortisol modulation of congruency effect* |  |  |  |  |  |  |  |  |
| *TP17 vs. TP14* |  |  |  |  |  |  |  |  |
| aPFC | L | 10 | 20 | -28 | 60 | -16 | <.0001 | 5.12 |
| Amygdala | R | 34 | 39 | 30 | -2 | -12 | <.0001 | 4.61 |
|  |  | 36 | 3 | 30 | 4 | -26 | .0003 | 3.62 |
| Pulvinar | R | - | 7 | 20 | -22 | 14 | .0002 | 3.70 |
| *TP20 vs. TP17* |  |  |  |  |  |  |  |  |
| aPFC | L | 10,11,47 | 575 | -26 | 54 | 2 | <.0001 | 5.64 |
|  | R | 11 | 116 | 20 | 54 | 0 | <.0001 | 3.56 |
| Amygdala | L | 34 | 47 | -18 | -4 | -14 | <.0001 | 5.63 |

Note: BA, Brodmann Area; k, number of voxels in a cluster; *p*, voxel-level significance value; *t*, t-statistic at the peak voxel; R, right; L, left; aPFC, anterior prefrontal cortex

#### Other related changes in the congruency effect

Irrespective of testosterone levels, there was a significant interaction between age and congruency in the aPFC between TP17 vs. TP14 (MNI coordinates: 24, 66, 2) and TP20 vs. TP17 (MNI coordinates: -28, 58, -12) suggesting a decrease in aPFC activity during emotion control compared to TP14 and TP20. The congruency effect was also significantly modulated by a testosterone x cortisol interaction for the TP17 vs. TP14 comparison in the aPFC, amygdala, and pulvinar and in the aPFC and amygdala for TP20 vs. TP17. These interaction terms were included in the model to control for mutual testosterone-cortisol effects, but are not described in detail as they go beyond the current scope and research question. For all ROI effects see Table 2; for whole-brain effects see Supplementary Information Table S1.

#### Functional connectivity

Having shown changes with age in aPFC and amygdala activation during emotional action control, we tested for testosterone-related changes in coupling between these two regions. Following previous studies (1, 2), we conducted psychophysiological interaction analyses during affect incongruent vs. affect congruent conditions of the AA task with the aPFC and amygdala as seed regions. We conducted the following ROI analyses: aPFC (seed) – amygdala (ROI) and amygdala (seed) – aPFC (ROI). There were no significant changes with age in connectivity strength as a function of testosterone. For remaining effects see Supplementary Information Table S2.

#### Behavioral Results and Hormone Levels

Testosterone values between the first and second sampling measurement (i.e. before (s1) and after (s2) the AA task) did not differ significantly at any of the three time-points either for males (TP14: *t*(19) = 0.99, *p* = 0.333; TP17: t(39) = 1.72, *p* = 0.093; TP20: *t*(35) = 1.47, p = 0.151) or females (TP14: *t*(25) = 0.66, *p* = 0.514; TP17: *t*(30) = 1.36, *p* = 0.184; TP20: *t*(31) = 1.85, *p* = 0.074). Testosterone s1 levels were used for all subsequent (behavioral and imaging) analyses in line with previous studies on the AA task (1, 2, 6).

A significant valence x response interaction for RTs confirmed AA task congruency effects as reported previously, driven by longer reaction times for affect-incongruent compared to affect-congruent responses (Figure 1D). The planned contrast for age revealed a significant overall decrease in RTs at TP17 compared to TP14 and a significant interaction for TP20 vs. TP17 with valence x testosterone x gender indicating that higher testosterone levels in 17-year old females were associated with faster RTs on happy trials (vs. angry), whereas for males higher testosterone levels were related to faster RTs on angry (vs. happy) trials at that age. However, the opposite association was present at age 20 where higher testosterone levels were associated with faster responding on angry trials for females and happy trials for males. Finally, there were significant main effects of response and affect, indicating slower RTs for avoid trials and angry faces. For all effects see Supplementary Information Table S3.

## Discussion

This longitudinal investigation tested the contribution of testosterone on prefrontal emotion control during the transition across adolescence and into young adulthood. In line with predictions from animal models, the positive effect of testosterone on aPFC engagement decreased from age 14 to age 17 and shifted by age 20, when higher testosterone levels were related to less aPFC activity. In contrast to adolescence, during young adulthood, testosterone - no longer related to pubertal development - impeded prefrontal control. In accordance with hallmarks of adult-like circuitry found in previous studies (2), this change in testosterone function is accompanied by increased testosterone-modulated amygdala reactivity during emotion control in young adults. Taken together, these findings point to a transition in the role of testosterone as a neurodevelopmental pubertal hormone and demonstrate the concurrent maturation of the prefrontal-amygdala circuit from middle adolescence to young adulthood.

It is well established that during adolescence, the prefrontal cortex undergoes structural and functional reorganization (49–51). Importantly, cognitive control enhances during emotional situations, which is accompanied by dynamic changes in prefrontal circuitry (52, 53) as well as cortico-cortico functional network strength, particularly with respect to pubertal development (54). The results of the present study extend these findings to cognitive control of automatic emotional actions and track it within individuals across three developmental time-points.

Rapid changes in testosterone levels during the first half of adolescence drive and organize neural development by modulating cell proliferation, cell death, and synaptic connectivity (4, 55–57). For example, previous neuroimaging studies in humans have implicated these organizational effects of testosterone to underlie prefrontal and subcortical structural changes across adolescence (20, 58–60). Such organizational effects decline by late adolescence, when testosterone starts to activate specific socio-sexual behaviors. The results of the current study translate findings from animal work on the organizational – activational hypothesis, specifically regarding context dependent social approach-avoidance motivation (15), by implementing a task designed to assess relevant social approach-avoidance behavior in humans. By doing so, we show that also within human individuals, the role of testosterone on aPFC emotion control transitions – first facilitating aPFC engagement during middle adolescence, to later impeding neural control by young adulthood.

Increased aPFC activity at age 14 replicates previous findings from this cohort (1) and is in line with other studies showing developmental shifts towards more prefrontal control in other domains of emotion-relevant processing (61, 62), including motor response inhibition (63, 64), reward sensitivity (8, 65, 66), and perceptual processing of emotional faces (67–70).

Diminished prefrontal control, coupled with increased amygdala reactivity in high-testosterone individuals has previously been shown in adults (2, 6) and parallels our findings at age 20. Crucially, however, within the same individuals - we demonstrated a switch from positive to negative prefrontal-testosterone modulation between ages 17 and 20. As such, the results of this study bridge seemingly conflicting findings from the developmental and adult literature. In line with these results and the notion of the changing role of testosterone across adolescence, (71) also showed similar trajectories in prefrontal and subcortical regions in response to emotional reactivity with respect to rising testosterone levels. The effect was specific in females, where a positive quadratic relationship in the amygdala and inverted U-shaped changes in other prefrontal regions (perigenual ACC) were observed. Finally, the present study extends previous findings of cumulative pubertal testosterone exposure on adult male human neural circuits (72) and supports the notion of testosterone-organizational effects in the human prefrontal cortex and amygdala.

The current findings place late adolescence at a pivotal point in the development of mature aPFC emotional regulation processes when both organizational and activational effects likely play a role. This is reflected in the different slope signs of the aPFC-testosterone effect at age 17, depending on the specific localization within the aPFC (and contrast). Given the region’s multifaceted roles as a domain-general center for monitoring alternative options (73) as well as coordinating dis-inhibition and enhancement processes in action selection circuits (74) – it is likely that the development of the aPFC itself may not be homogenous at this time-point.

Contrary to expectations, we did not find changes in aPFC-amygdala functional connectivity with respect to testosterone levels. Given the role of testosterone in driving both functional and structural development, perhaps task-related developmental connectivity changes are better reflected in the latter. For example, a multidimensional structural brain development index has been shown to accurately delineate developmental trajectories related to cognitive performance (75). Specifically for the AA task, social-emotional control abilities in adults have been shown to depend on the strength of structural connectivity between the aPFC and amygdala (76).

The testosterone-modulated emergence of an adult-like aPFC-amygdala circuit for regulating social emotional actions likely involves more complex interactions with other hormones, such as cortisol. The interplay between testosterone and cortisol is likely related to maturational processes due to the integrated synthesis and metabolism of cortisol and sex hormones by the HPA and HPG axes (77). It can, therefore, be expected that mutual testosterone-cortisol effects also undergo a transition across adolescence (78–80). In our study, this is evidenced by combined modulation of testosterone and cortisol of the aPFC and amygdala across all three time-points.

### Interpretational issues

The present study addresses the transition in organization-activational effects of testosterone on aPFC control assuming that this is not a dichotomous change, but most likely a gradual shift between organizational and activational effects. The exact mechanism by which neural organizational effects close is not clear. Opening and closing of sensitivity periods and associated hormone-dependent neural plasticity is known to involve changes in neurotransmitter and neurotrophic factor expression, dendritic morphology (e.g., spine formation, pruning), myelination, and regulation of cell numbers (81). For example, normative decreases of frontal cortex plasticity may be related to changes in BDNF expression driven by pubertal testosterone (82). Perhaps the transition to testosterone-activational effects is a result of reorganization or consolidation at the circuit and neuronal level, which also limits the capacity for further testosterone-dependent organization (83).

The choice of testosterone as an objective and physiological marker of pubertal development in both females and males is supported by previous work (84, 85) and the organizational effects of this hormone on neural development during adolescence (5, 59, 86). Because testosterone can act on receptors through its conversion to other steroid hormones, it can have similar effects in women and men (87). Importantly, the testosterone effects reported here reflect within-sex variation: testosterone levels were standardized within sex, so its effects cannot be attributed to relative differences in hormonal levels between females and males. Future studies should test whether the effects in females obtained outside the menstrual phase replicate when taking into account the precise phase of the menstrual cycle. Similarly, future studies should replicate these findings in a larger dataset that enables modeling multiple interactions with testosterone, cortisol, and sex that may underlie more complex developmental patterns.

When comparing testosterone effects at TP17 versus those at TP14, the strongest activation was in the left aPFC, whereas for TP17 versus TP20 in the right aPFC. Importantly, the same direction of effects was present for both comparisons also in the contralateral hemisphere at subthreshold levels (see Figure 1B). It is, therefore, likely that the effect of testosterone on prefrontal control may be strongest in specific localizations within the aPFC, reflecting the heterogenous development of this region.

The behavioral regulation of social emotional actions was replicated over all time-points, but was not coupled with aPFC – testosterone changes across age. However, the AA task used in this study was designed to evoke a mild social challenge that, in a healthy cohort, does not lead to behavioral differences. This feature of the experimental design was selected to isolate pubertal maturational effects at the neural level, avoiding interpretational confounds due to differences in task performance (e.g., related to differences in skill acquisition or puberty-related reorganization of behavioral repertoire; for review see (88)).

The current study did not investigate whether individual increases or decreases in testosterone levels correspond to changes in neural activity, but rather whether the *function* of testosterone changed across adolescence and adulthood. Although relevant, this research question goes beyond the scope of this paper.

## Conclusion

With the use of a prospective longitudinal design, the present study sought to characterize the functional shift in testosterone from middle to late adolescence and its association with the neural control of emotional actions. Our findings support a decrease in the organizational effects of testosterone in the aPFC and the concurrent transition into activational effects by young adulthood. These results are relevant for neuro-endocrine models of pubertal development by explicating the mechanisms involved in the maturation of testosterone-modulated emotional action control. Chronic failures in controlling emotional action tendencies are common in various affective disorders, whose onset peaks in adolescence (89). This study opens the possibility of testing whether altered neuro-endocrine trajectories are mechanistically related to the development of psychopathology, such as social anxiety and aggression.

## Acknowledgements

The authors would like to thank all the families participating in the Nijmegen Longitudinal Study who made this research possible..

