## Supplementary Information for "Developmental shift in testosterone influence on prefrontal emotion control"

**Table S1.**

Whole-brain significant clusters and peak activation changes with age during affect incongruent compared to affect congruent trials in the AA task

| Anatomical region | Side | BA | k | MNI coordinates x | y | z | *p* | t |
|---|---|---|---|---|---|---|---|---|
| *Congruency effect* | | | | | | | | |
| *TP17 vs. TP14* | | | | | | | | |
| Anterior cingulate cortex/ Olfactory gyrus | L | 25 | 1103 | -2 | 24 | -2 | <.0001 | 4.52 |
| Inferior occipital gyrus | R | 18,19 | 676 | 34 | -80 | -8 | <.0001 | 4.57 |
| Superior temporal / postcentral gyrus | L | 43,48 | 534 | -66 | -4 | 6 | <.0001 | 4.65 |
| Postcentral gyrus | L | 3,41 | 379 | -38 | -40 | 28 | <.0001 | 4.55 |
| *TP20 vs. TP17* | | | | | | | | |
| Lingual / calcarine gyrus | L | 18,19 | 893 | -16 | -78 | -2 | <.0001 | 4.44 |
| Postcentral gyrus | L | 43 | 505 | -66 | 0 | 18 | <.0001 | 4.38 |
| *Testosterone modulation of congruency effect* | | | | | | | | |
| *TP17 vs. TP14* | | | | | | | | |
| Insula | R | 48 | 958 | 32 | 10 | -4 | <.0001 | 4.54 |
| Gyrus rectus / SFG | R | 11,25 | 516 | 6 | 12 | -26 | <.0001 | 4.91 |
| *TP20 vs. TP17* | | | | | | | | |
| Supramarginal gyrus | L | 48 | 772 | -52 | -42 | 30 | <.0001 | 4.48 |
| Inferior parietal gyrus | R | 39,40 | 460 | 48 | -56 | 42 | <.0001 | 4.53 |
| Cerebellum | R | - | 370 | -14 | -74 | -30 | .0007 | 3.42 |
| *Testosterone and cortisol modulation of congruency effect* | | | | | | | | |
| *TP17 vs. TP14* | | | | | | | | |
| Insula / Putamen | R | 47,48 | 849 | 24 | 24 | -2 | <.0001 | 4.56 |
| Cuneus | R | 18 | 568 | 4 | -90 | 24 | <.0001 | 4.56 |
| Gyrus rectus | R/L | 11 | 406 | 0 | 38 | -28 | <.0001 | 7.29 |
| SFG / middle frontal / WM | R | 6,8 | 379 | 24 | 4 | 32 | <.0001 | 4.41 |

| | | | | | | | | |
|---|---|---|---|---|---|---|---|---|
| *TP20 vs. TP17* | | | | | | | | |
| Gyrua rectus | R/L | 11 | 1934 | 0 | 36 | -28 | <.0001 | 5.13 |
| Angular / inferior parietal gyrus | L | 39,40 | 1127 | -50 | -54 | 30 | <.0001 | 4.50 |
| Postcentral / precentral gyrus | L | 4,43 | 1054 | -66 | -8 | 24 | <.0001 | 4.40 |
| Middle frontal | L | 9,46 | 622 | -38 | 30 | 42 | <.0001 | 4.67 |
| Precentral gyrus | R | 6 | 361 | 50 | -2 | 50 | <.0001 | 4.38 |

Note: BA, Brodmann Area; k, number of voxels in a cluster; *p*, voxel-level significance value; t, t-statistic at the peak voxel; R, right; L, left. SFG, superior frontal gyrus

**Table S2.**

Functional connectivity between amygdala (seed, MNI coordinates: 20, 0, -14) and aPFC

| Anatomical region | Side | BA | k | MNI coordinates | | | *p* | t |
|---|---|---|---|---|---|---|---|---|
| | | | | x | y | z | | |
| *TP20 vs. TP17* | | | | | | | | |
| aPFC | L | 10 | 15 | -42 | 60 | -6 | <.0001 | 4.57 |
| *TP17 vs. TP14 testosterone and cortisol modulation* | | | | | | | | |
| aPFC | L | 47 | 6 | -38 | 52 | -12 | <.0001 | 4.17 |

Note: BA, Brodmann Area; k, number of voxels in a cluster; *p*, voxel-level significance value; t, t-statistic at the peak voxel; R, right; L, left; aPFC, antererior prefrontal cortex

**Table S3.**
AA task - behavioral model statistics for the change in the congruency effect with age and in association with testosterone, cortisol, and gender

| Fixed effects | | | | |
|---|---|---|---|---|
| | **Estimate** | **Est. Error** | **95% IC** | |
| Intercept | -0.55 | 0.02 | -0.59, -0.52 | |
| Response | 0.04 | 0.01 | 0.03, 0.05 | * |
| Valence | 0.03 | 0.01 | 0.01, 0.05 | * |
| Testosterone | -0.03 | 0.01 | -0.05, 0 | |
| Cortisol | 0 | 0.01 | -0.02, 0.02 | |
| Gender | -0.05 | 0.03 | -0.11, 0.01 | |
| Valence X Response | -0.07 | 0.01 | -0.09, -0.05 | * |
| Response X Testosterone | 0.01 | 0.01 | 0, 0.02 | |
| Valence X Testosterone | 0 | 0 | -0.01, 0 | |
| Response X Cortisol | -0.01 | 0.01 | -0.02, 0 | |
| Valence X Cortisol | 0 | 0 | -0.01, 0 | |
| Response X Gender | -0.01 | 0.01 | -0.03, 0.02 | |
| Valence X Gender | 0.01 | 0.01 | -0.01, 0.02 | |
| Testosterone X Gender | 0.01 | 0.02 | -0.04, 0.06 | |
| Cortisol X Gender | -0.02 | 0.02 | -0.06, 0.02 | |
| Testosterone X Cortisol | 0 | 0.01 | -0.02, 0.03 | |
| Valence X Response X Testosterone | 0 | 0.01 | -0.02, 0.02 | |
| Valence X Response X Cortisol | 0.01 | 0.01 | -0.01, 0.03 | |
| Valence X Response X Gender | 0.01 | 0.02 | -0.03, 0.05 | |
| Valence X Testosterone X Cortisol | 0.01 | 0 | 0, 0.02 | |
| Response X Testosterone X Cortisol | 0 | 0.01 | -0.01, 0.01 | |
| Valence X Testosterone X Gender | 0 | 0.01 | -0.02, 0.01 | |
| Response X Testosterone X Gender | 0.01 | 0.01 | -0.02, 0.03 | |
| Valence X Cortisol X Gender | -0.01 | 0.01 | -0.02, 0.01 | |
| Response X Cortisol X Gender | -0.02 | 0.01 | -0.04, 0 | |
| Testosterone X Cortisol X Gender | -0.02 | 0.02 | -0.07, 0.02 | |
| Valence X Response X Testosterone X Cortisol | -0.01 | 0.01 | -0.03, 0.01 | |
| Valence X Response X Testosterone X Gender | 0 | 0.02 | -0.04, 0.04 | |
| Valence X Response X Cortisol X Gender | -0.01 | 0.02 | -0.05, 0.02 | |
| Valence X Testosterone X Cortisol X Gender | 0 | 0.01 | -0.02, 0.01 | |
| Response X Testosterone X Cortisol X Gender | 0 | 0.01 | -0.02, 0.02 | |
| Valence X Response X Testosterone X Cortisol X Gender | 0 | 0.02 | -0.04, 0.04 | |

| | | | | |
|---|---|---|---|---|
| *Contrast TP17 - TP14* | | | | |
| Age | -0.11 | 0.02 | -0.15, -0.07 | * |
| Age X Valence | -0.01 | 0.01 | -0.03, 0.01 | |
| Age X Response | 0.01 | 0.01 | -0.01, 0.03 | |
| Age X Testosterone | 0.04 | 0.03 | -0.02, 0.01 | |
| Age X Cortisol | -0.01 | 0.02 | -0.05, 0.04 | |
| Age X Gender | -0.03 | 0.04 | -0.11, 0.05 | |
| Age X Valence X Response | 0 | 0.02 | -0.04, 0.04 | |
| Age X Response X Testosterone | -0.02 | 0.01 | -0.05, 0.01 | |
| Age X Valence X Testosterone | 0.02 | 0.01 | 0, 0.04 | |
| Age X Response X Cortisol | 0.01 | 0.01 | -0.01, 0.04 | |
| Age X Valence X Cortisol | 0 | 0.01 | -0.02, 0.02 | |
| Age X Response X Gender | 0.02 | 0.02 | -0.02, 0.06 | |
| Age X Valence X Gender | 0.01 | 0.02 | -0.02, 0.05 | |
| Age X Testosterone X Gender | -0.07 | 0.06 | -0.18, 0.04 | |
| Age X Cortisol X Gender | 0.08 | 0.05 | -0.02, 0.19 | |
| Age X Testosterone X Cortisol | -0.01 | 0.03 | -0.08, 0.05 | |
| Age X Valence X Response X Testosterone | 0 | 0.02 | -0.05, 0.05 | |
| Age X Valence X Response X Cortisol | 0.01 | 0.02 | -0.03, 0.05 | |
| Age X Valence X Response X Gender | 0.05 | 0.04 | -0.02, 0.13 | |
| Age X Valence X Testosterone X Cortisol | -0.01 | 0.01 | -0.04, 0.01 | |
| Age X Response X Testosterone X Cortisol | -0.02 | 0.01 | -0.05, 0 | |
| Age X Valence X Testosterone X Gender | 0.03 | 0.02 | -0.01, 0.07 | |
| Age X Response X Testosterone X Gender | -0.01 | 0.03 | -0.06, 0.04 | |
| Age X Valence X Cortisol X Gender | -0.01 | 0.02 | -0.05, 0.03 | |
| Age X Response X Cortisol X Gender | 0.01 | 0.02 | -0.04, 0.05 | |
| Age X Testosterone X Cortisol X Gender | -0.1 | 0.06 | -0.22, 0.02 | |
| Age X Valence X Response X Testosterone X Cortisol | -0.03 | 0.03 | -0.08, 0.02 | |
| Age X Valence X Response X Testosterone X Gender | 0.09 | 0.05 | 0, 0.18 | |
| Age X Valence X Response X Cortisol X Gender | -0.06 | 0.04 | -0.14, 0.03 | |
| Age X Valence X Testosterone X Cortisol X Gender | 0.01 | 0.02 | -0.03, 0.05 | |
| Age X Response X Testosterone X Cortisol X Gender | -0.03 | 0.03 | -0.08, 0.03 | |
| Age X Valence X Response X Testosterone X Cortisol X Gender | 0 | 0.05 | -0.1, 0.1 | |
| *Contrast TP20 - TP17* | | | | |
| Age | -0.02 | 0.02 | -0.06, 0.02 | |
| Age X Valence | -0.01 | 0.01 | -0.02, 0.01 | |
| Age X Response | -0.01 | 0.01 | -0.03, 0.01 | |

| | | | | |
|---|---|---|---|---|
| Age X Testosterone | 0 | 0.02 | -0.05, 0.05 | |
| Age X Cortisol | -0.01 | 0.02 | -0.05, 0.03 | |
| Age X Gender | 0.02 | 0.04 | -0.06, 0.09 | |
| Age X Valence X Response | -0.03 | 0.02 | -0.07, 0 | |
| Age X Response X Testosterone | 0 | 0.01 | -0.02, 0.02 | |
| Age X Valence X Testosterone | -0.01 | 0.01 | -0.02, 0.01 | |
| Age X Response X Cortisol | -0.01 | 0.01 | -0.03, 0.01 | |
| Age X Valence X Cortisol | -0.01 | 0.01 | -0.02, 0.01 | |
| Age X Response X Gender | 0 | 0.02 | -0.04, 0.03 | |
| Age X Valence X Gender | -0.02 | 0.01 | -0.05, 0.01 | |
| Age X Testosterone X Gender | -0.06 | 0.05 | -0.15, 0.04 | |
| Age X Cortisol X Gender | 0.04 | 0.05 | -0.05, 0.13 | |
| Age X Testosterone X Cortisol | 0.01 | 0.03 | -0.04, 0.07 | |
| Age X Valence X Response X Testosterone | 0 | 0.02 | -0.04, 0.05 | |
| Age X Valence X Response X Cortisol | -0.01 | 0.02 | -0.04, 0.03 | |
| Age X Valence X Response X Gender | 0 | 0.03 | -0.06, 0.07 | |
| Age X Valence X Testosterone X Cortisol | -0.01 | 0.01 | -0.02, 0.01 | |
| Age X Response X Testosterone X Cortisol | 0 | 0.01 | -0.03, 0.02 | |
| Age X Valence X Testosterone X Gender | -0.05 | 0.02 | -0.09, -0.01 | * |
| Age X Response X Testosterone X Gender | 0.01 | 0.02 | -0.03, 0.06 | |
| Age X Valence X Cortisol X Gender | -0.01 | 0.02 | -0.04, 0.03 | |
| Age X Response X Cortisol X Gender | -0.01 | 0.02 | -0.05, 0.03 | |
| Age X Testosterone X Cortisol X Gender | 0.01 | 0.05 | -0.1, 0.11 | |
| Age X Valence X Response X Testosterone X Cortisol | 0.03 | 0.02 | -0.01, 0.07 | |
| Age X Valence X Response X Testosterone X Gender | -0.06 | 0.04 | -0.13, 0.02 | |
| Age X Valence X Response X Cortisol X Gender | 0.05 | 0.04 | -0.03, 0.12 | |
| Age X Valence X Testosterone X Cortisol X Gender | -0.03 | 0.02 | -0.07, 0.01 | |
| Age X Response X Testosterone X Cortisol X Gender | -0.02 | 0.02 | -0.07, 0.02 | |
| Age X Valence X Response X Testosterone X Cortisol X Gender | 0 | 0.04 | -0.09, 0.08 | |
